# ClpK as a Chaperone Hub: Computational Exploration of its Protein Interaction Landscape

**DOI:** 10.1101/2025.11.28.691151

**Authors:** Tehrim Ballim, Ikechukwu Achilonu, Thandeka Khoza

## Abstract

Protein-protein interaction networks provide a foundation for understanding the molecular mechanisms within cells and are increasingly recognized as promising targets for therapeutic intervention. However, the experimental elucidation of these interactions is complicated by the complexity and dynamic nature of protein interaction networks. *In silico* methods have become invaluable allowing researchers to identify potential interactions and focus on those that may be biologically significant to guide experimental validation and drug discovery. This study employed computational modelling, visualization, and network analysis to investigate the interactions of ClpK, a Clp ATPase integral to protein homeostasis and thermotolerance of *Klebsiella pneumoniae*. Potential interacting partners of ClpK were predicted using the STRING database with ClpB used as a structural proxy for ClpK. The proteins with edge confidence scores greater than 0.8 were selected for molecular docking and molecular dynamics simulations to assess interaction stability and binding affinities. Molecular docking using the ClusPro server confirmed stable interactions between ClpK and its predicted partners, which were further characterised using PDBsum and Hawkdock to visualise binding interfaces and estimate binding free energies MM-GBSA calculation. Among the predicted complexes, KPN_01323 had the highest interaction score, highlighting it as the most promising candidate for further investigation. Molecular dynamics simulations further confirmed the stabilisation of putative partners upon interaction with ClpK supporting the robustness of this interaction. Collectively, these findings provide insight into the ClpK interaction network and establish a framework for future *in vitro* and *in vivo* studies contributing to elucidate the biological role and therapeutic potential of ClpK.

## 1.1. Introduction

Bacteria depend on an ATP dependent protease system to regulate protein turnover and eliminate misfolded or dysfunctional proteins, particularly under environmental stress conditions (Queraltó *et al*., 2023, Xu *et al*., 2024). The Clp/Hsp100 family of molecular chaperones forms a key component of this system, functioning in both protein disaggregation and targeted proteolysis under stressful conditions (Maurizi and Li, 2001). This family consists of Clp ATPase and the caseinolytic proteolytic subunit (ClpP), which together assemble into the caseinolytic protein (Clp) complex responsible for ATP-dependent protein degradation (Bouchnak and Van Wijk, 2021, Brötz-Oesterhelt and Sass, 2014). Clp ATPases are classified into two classes; Class I ATPase which contain two ATP binding domains separated by a spacer region (e.g.: ClpA, ClpB, ClpK, ClpC) and Class II ATPases which contain a single ATP-binding domain (e.g.: ClpY and ClpX) (Motiwala *et al*., 2021, Queraltó *et al*., 2023, Yu *et al*., 2018).

ClpK belongs to Class I ATPase and has been associated with heat stress survival in *K. pneumoniae* (Bojer *et al*., 2010, Bojer *et al*., 2013, Motiwala *et al*., 2022). Interestingly, ClpK is considered a unique Clp ATPase among Gram-negative bacteria because it contains an intermediate-size linker domain resembling that of ClpC and its structural homologs, ClpE and ClpL (Bojer *et al*., 2010). This feature suggests that ClpK may employ specialized molecular interactions that differentiate its functional behaviour from other Clp homologs. Consistent with this observation, adaptor proteins such as MecA and ClpS have been shown to interact with other Clp ATPases to enhance proteolysis and modulate substrate recognition (Wahl *et al*., 2014, Dougan *et al*., 2002). In addition to Clp ATPase-adaptor interaction, chaperone activity is also critical for maintaining protein quality control. For example, the molecular chaperone DnaK interacts with ClpB under stress conditions to deliver misfolded substrates to the proteolytic complex for degradation (Schlieker *et al*., 2004). Therefore, such proteinprotein interactions are essential for maintaining bacterial proteostasis, survival, stress tolerance, and virulence in pathogens such as *Staphylococcus aureus* and *Mycobacterium tuberculosis*, where Clp systems have been identified as promising drug targets (Ollinger *et al*., 2012, Frees *et al*., 2014).

Among these systems, Clp ATPases such as ClpA, ClpX and ClpC contain the ClpP interaction motif – IG(F/L) which mediates their association with the ClpP peptidase to form a proteolytic complex responsible for protein degradation (Maurizi and Xia, 2004, Queraltó *et al*., 2023). While these interactions are well characterized, the interaction partners and structural basis of ClpK-mediated associations with adaptor proteins or chaperones remains unexplored. Furthermore, no crystal structures, cryo-electron microscopy (cryo-EM) data, or experimentally validated interaction models for *K. pneumoniae* ClpK are available in structural databases. This lack of structural and functional insight represents a major gap in understanding how ClpK mediates substrate recognition, binding and protein degradation during cellular stress. Addressing this gap is critical for exploiting ClpK as a potential antimicrobial target, especially in the context of rising multidrug resistance in *K. pneumoniae*. Elucidating these interactions is crucial for defining ClpK’s cellular role and aid in the rational design of inhibitors that disrupt key chaperone-protease interactions.

Given the importance of PPIs in regulating Clp ATPase function, mapping the interaction network of ClpK is essential for understanding its biological mechanisms. Computational PPI prediction tools such as Search Tool for the Retrieval of Interacting Genes/Proteins (STRING) and deep-learning models now enable the identification of potential interaction partners, complementing experimental approaches such as coimmunoprecipitation or X-ray crystallography (Farooq *et al*., 2021). The STRING database integrates both known and predicted PPIs from diverse sources, including experimental data, computational prediction methods, and public text collections (Szklarczyk *et al*., 2023, Ding and Kihara, 2018). It then assigns interaction scores based on evidence, allowing for the identification of high-confidence interaction partners for a given protein (Szklarczyk *et al*., 2023). To explore the structural basis of these interactions, protein structure prediction platforms such as AlphaFold and SWISS-MODEL are then widely used (Jumper *et al*., 2021, Schwede *et al*., 2003). AlphaFold uses deep learning to generate high-accuracy 3D models of proteins based on their amino acid sequences, while SWISS-MODEL uses homology-based modelling for reliable structure prediction in the absence of crystal structures (Jumper *et al*., 2021, Waterhouse *et al*., 2024). To predict and analyse protein–protein binding modes, molecular docking tools such as ClusPro and HADDOCK are used. These tools simulate interactions between two or more protein structures by considering complementarity in shape, electrostatics, and known binding residues, thereby predicting plausible interaction interfaces and binding affinities (De Vries *et al*., 2010, Kozakov *et al*., 2017).

In this study the ClpK interaction network was first investigated using the STRING database, subsequently, homology models of ClpK and its putative interaction partners were modelled successfully for structural and functional analysis. These models were then subjected to docking and molecular dynamics simulations, which provided insights into the structural basis and stability of these interactions, laying the groundwork for future experimental validation and inhibitor development.

## 1.2. Results

### 1.2.1. Identification of ClpK Interacting Proteins

Interaction networks were constructed from the STRING database using ClpB from *Klebsiella pneumoniae* MGH78578 as a proxy for ClpK due to its highest sequence identity (44.3%; Figure 1A) amongst other homologs. A high-confidence homology-based association was inferred with an Expect value (E-value) of 7.5×10^−188^, suggesting strong evolutionary conservation and supporting the reliability of predicted ClpK interactions. The resulting interaction network revealed ClpB exhibits dense connectivity with multiple chaperones and protease-related proteins (Figure 1B). Notably, strong associations were predicted with molecular chaperones DnaK, GrpE, ClpP, and DnaJ, all of which are integral components of the cellular protease machinery. The edge confidence scores of these interactions ranged from 0.839 to 0.976, suggesting evidence for functional linkage (Szklarczyk *et al*., 2023). It is important to note that STRING network edges represent protein–protein associations, signifying a shared or related biological function. These associations do not necessarily imply direct physical binding but may reflect participation in the same biological pathway or process (Szklarczyk *et al*., 2023).

**Figure 1:**
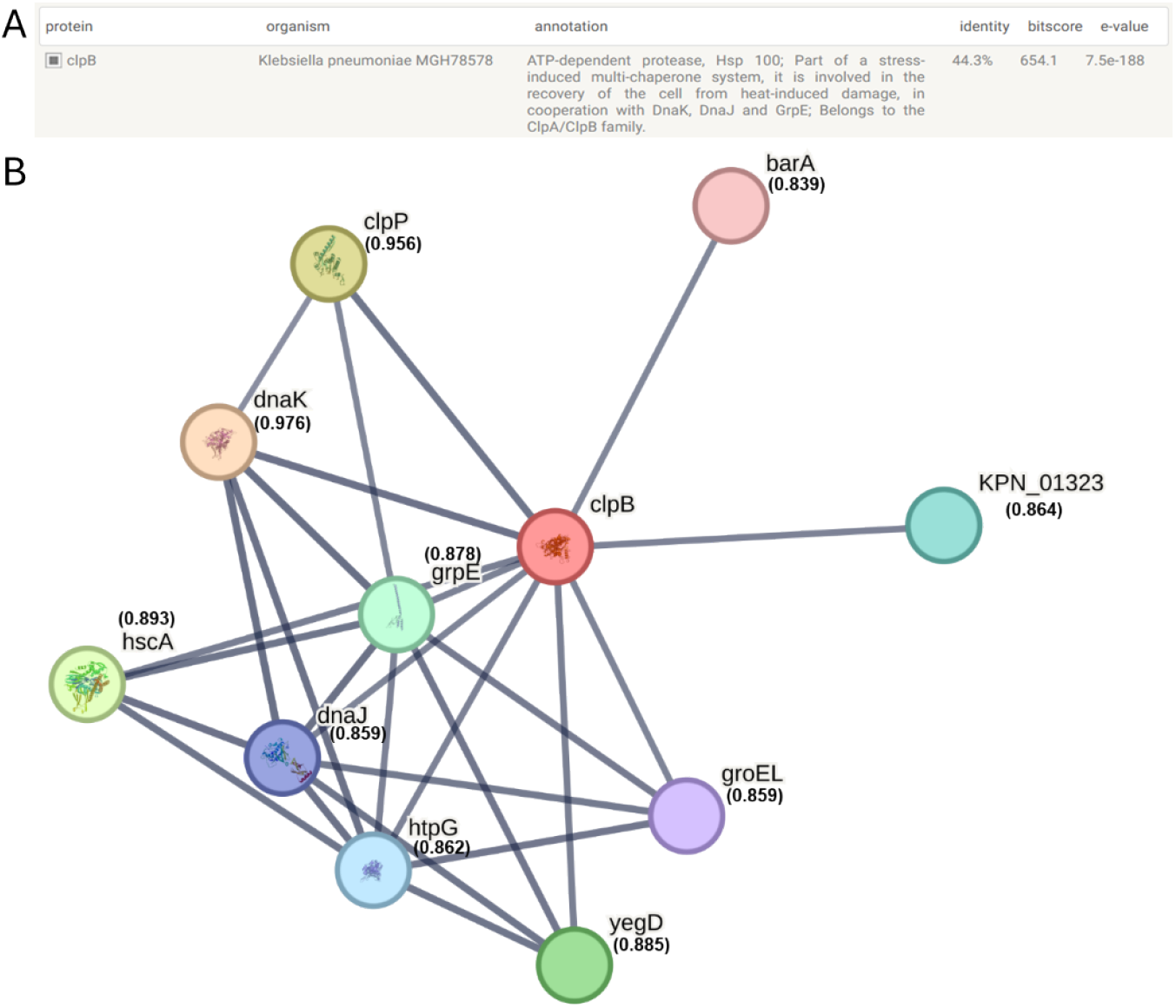
Predicted protein-protein interaction network of ClpK. **A)** *Klebsiella pneumoniae* MGH78578 ClpB was used as a proxy for ClpK due to its highest sequence identity. **B)** The network depicts predicted ClpK protein-protein interactions, identified using the STRING database. Each node represents a protein, and edges indicate functional associations based on experimental data, text mining, co-expression, and computational prediction. The key interacting proteins are *clpP* (Uniprot ID: A6T5I0), *dnaK* (Uniprot ID: A6T4F4), *dnaJ* (Uniprot ID: A6T4F5), *grpE* (Uniprot ID: A6TCM1), *htpG* (Uniprot ID: A6T5N6), *groEL* (Uniprot ID: A6TH28), *hscA* (Uniprot ID: A6TCE7), *barA* (Uniprot ID: A6TD58), *KPN_01323* (Uniprot ID: A6T834), and *yegD* (Uniprot ID: A6TBG8). The edge confidence scores are black, bold and in brackets.

### 1.2.2. Modelling of ClpK and its putative interacting partners

Structural models of ClpK and its putative interacting partners were generated using the SwissModel server (Figure 2A). Model quality was successfully evaluated prior to multiple validation metrics analysis using the Molprobity score, percentage of residues in Ramachandran favoured regions and QMEANDisCo global scores (Figure 2B). The MolProbity score is generated using a composite measure in which lower values correspond to models of superior structural quality (Chen *et al*., 2010). The clash score, rotamer and Ramachandran evaluations were combined into a single composite metric, where lower scores indicate higher structural quality (Chen *et al*., 2015). For all the modelled proteins, more than 80% of the residues fell within the favoured regions of the Ramachandran plot. Additionally, the QMEANDisCo global score, which ranges from 0 to 1 and reflects both global and per-residue model quality, was close to 1 for all the models, indicating high structural reliability (Studer *et al*., 2019).

**Figure 2:**
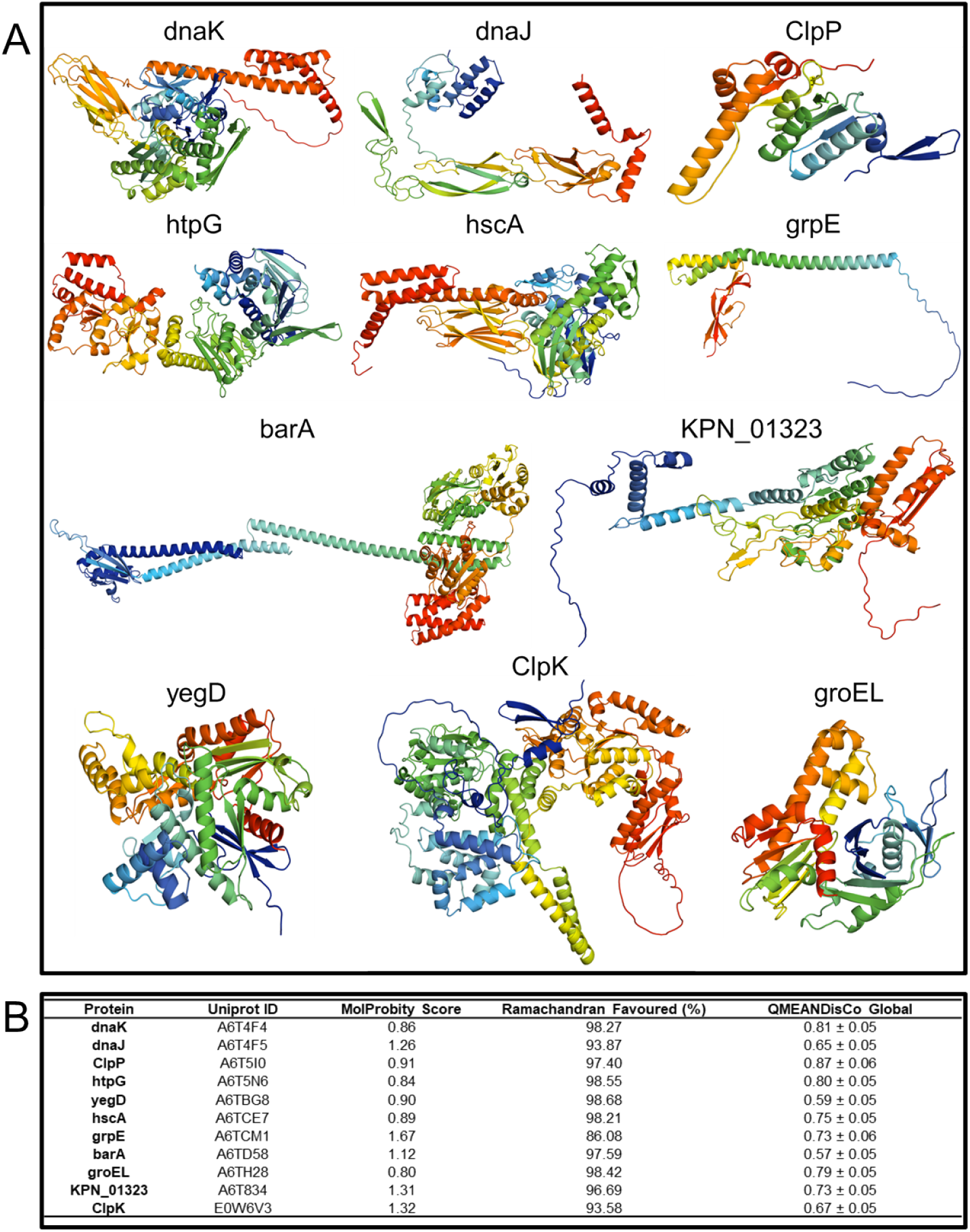
Homology modelling of ClpK and predicted interacting partners. **A)** The models were generated using the SwissModel server and visualised using PyMol (Schrödinger, 2015, Schwede *et al*., 2003). **B)** Summary of the structural validation parameters for the modelled proteins.

### 1.2.3. Docking analysis of ClpK with putative interaction partners

Interactions between ClpK and its predicted interacting partners were analysed through rigid-body protein-protein docking using the ClusPro 2.0 server. This server uses the blind rigid PIPER docking algorithm based on fast Fourier transformation (FFT) correlation to efficiently sample low-energy conformations using pairwise interaction potentials (Kozakov *et al*., 2017). Top-ranking blind-docking models (Table 1) were selected based on cluster sizes and energy parameters, with larger clusters and lower weighted scores representing the most probable binding conformations. Among the predicted complexes, ClpP, dnaJ and barA had the lowest energy binding scores (-1341.3, -1396.5 and -1565.6, respectively), indicating a strong likelihood of stable interaction with ClpK.

**Table 1:** Blind docking showing models with the highest cluster size and lowest weighted scores.

| Protein | Members | Representative | Weighted Score |
| --- | --- | --- | --- |
| DnaK | 31 | Center | -701.7 |
|  |  | Lowest Energy | -810.5 |
| GrpE | 35 | Center | -934.1 |
|  |  | Lowest Energy | -997.0 |
| groEL | 36 | Center | -818.7 |
|  |  | Lowest Energy | -885.0 |
| hscA | 37 | Center | -930.3 |
|  |  | Lowest Energy | -1019.6 |
| yegD | 46 | Center | -1015.3 |
|  |  | Lowest Energy | -1015.3 |
| KPN_01323 | 48 | Center | -1215.5 |
|  |  | Lowest Energy | -1223.0 |
| barA | 53 | Center | -1467.4 |
|  |  | Lowest Energy | -1565.6 |
| dnaJ | 69 | Center | -1358.0 |
|  |  | Lowest Energy | -1396.5 |
| htpG | 59 | Center | -1106.6 |
|  |  | Lowest Energy | -1106.6 |
| ClpP | 100 | Center | -1134.4 |
|  |  | Lowest Energy | -1341.3 |

The docking poses revealed variable binding orientations and interface regions, suggesting different potential interaction modes for each partner (Figure 3). It was observed that the putative partners bind near of the N-terminal domain, except hscA and groEL which bind in the proximity of the C-terminal domain. These predicted complexes suggest a range of potential binding modes suggesting functional diversity among the putative interactors.

**Figure 3:**
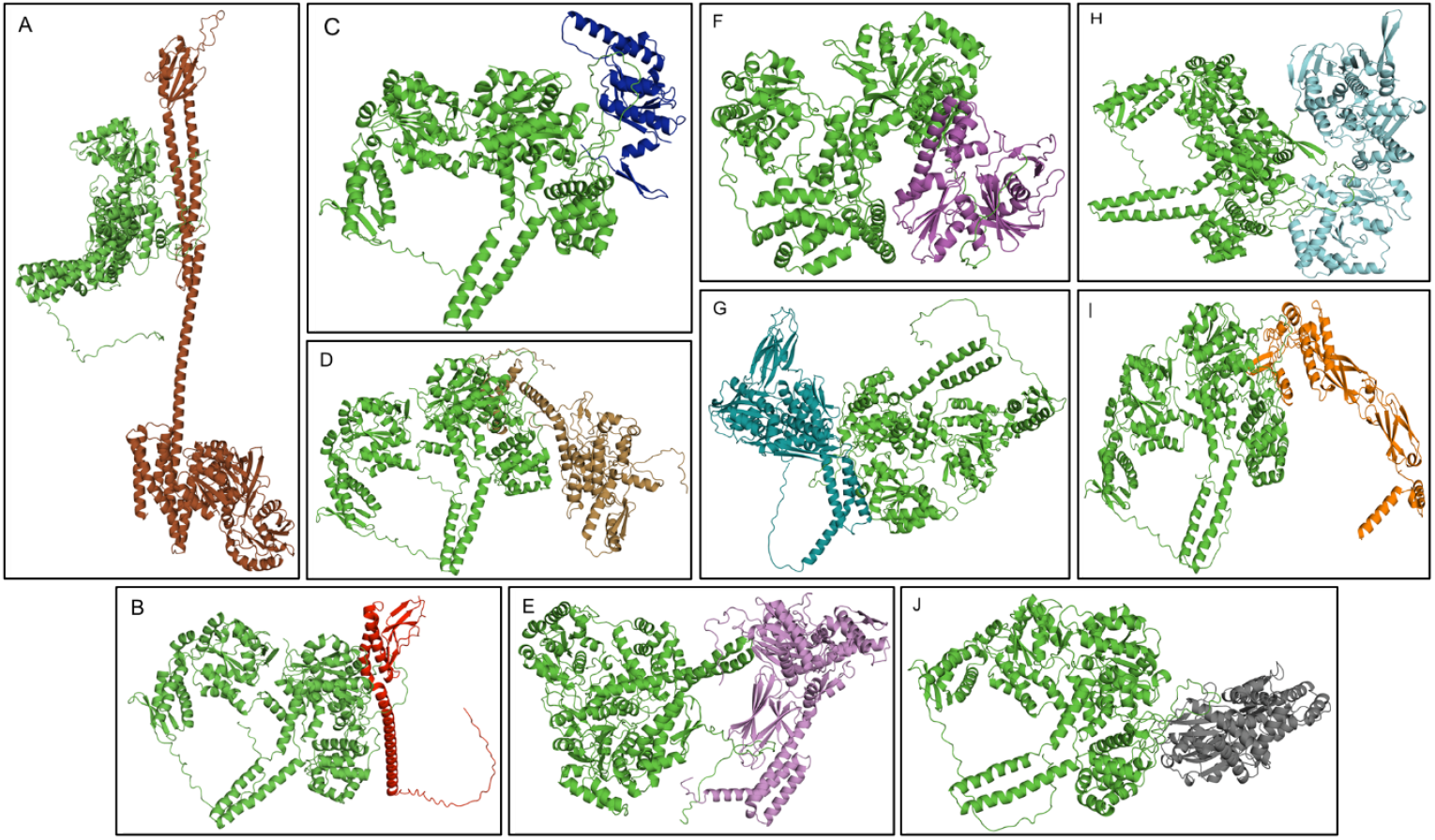
ClpK-putative partner docking complexes. ClpK was coloured green and is presented docked with **A)** barA (brown), **B)** grpE (red), **C)** ClpP (blue), **D)** KPN_01323 (light brown), **E)** hscA (violet), **F)** groEL (purple), **G)** DnaK (teal), **H)** htpG (light blue), **I)** DnaJ (orange), and **J)** YegD (grey). The ClpK complexes were visualised using PyMol.

Analysis of predicted interaction interfaces identified key residues mediating the stability of ClpK-partner complexes (Figure 4). The complexes were primarily stabilised by a combination of salt bridges, hydrogen bonds, and non-bonded contacts, which collectively contribute to protein–protein interaction specificity, stability and recognition (Donald *et al*., 2011). As shown in Figure 4A and Figure 4B, KPN_01323 exhibited the highest number of intermolecular interactions, whereas DnaK displayed the fewest. This trend is consistent with the binding energy scores presented in Table 1 where DnaK had the weakest predicted binding affinity and KPN_01323 was among the proteins with the strongest interaction energies. KPN_01323’s high interaction count highlights its potential as physiologically relevant binding partner of ClpK. These findings provide structural rationale to prioritise ClpK-KPN_01323 as a candidate for future *in silico* studies, especially in the context of ClpK’s role in stress response and protein homeostasis.

**Figure 4:**
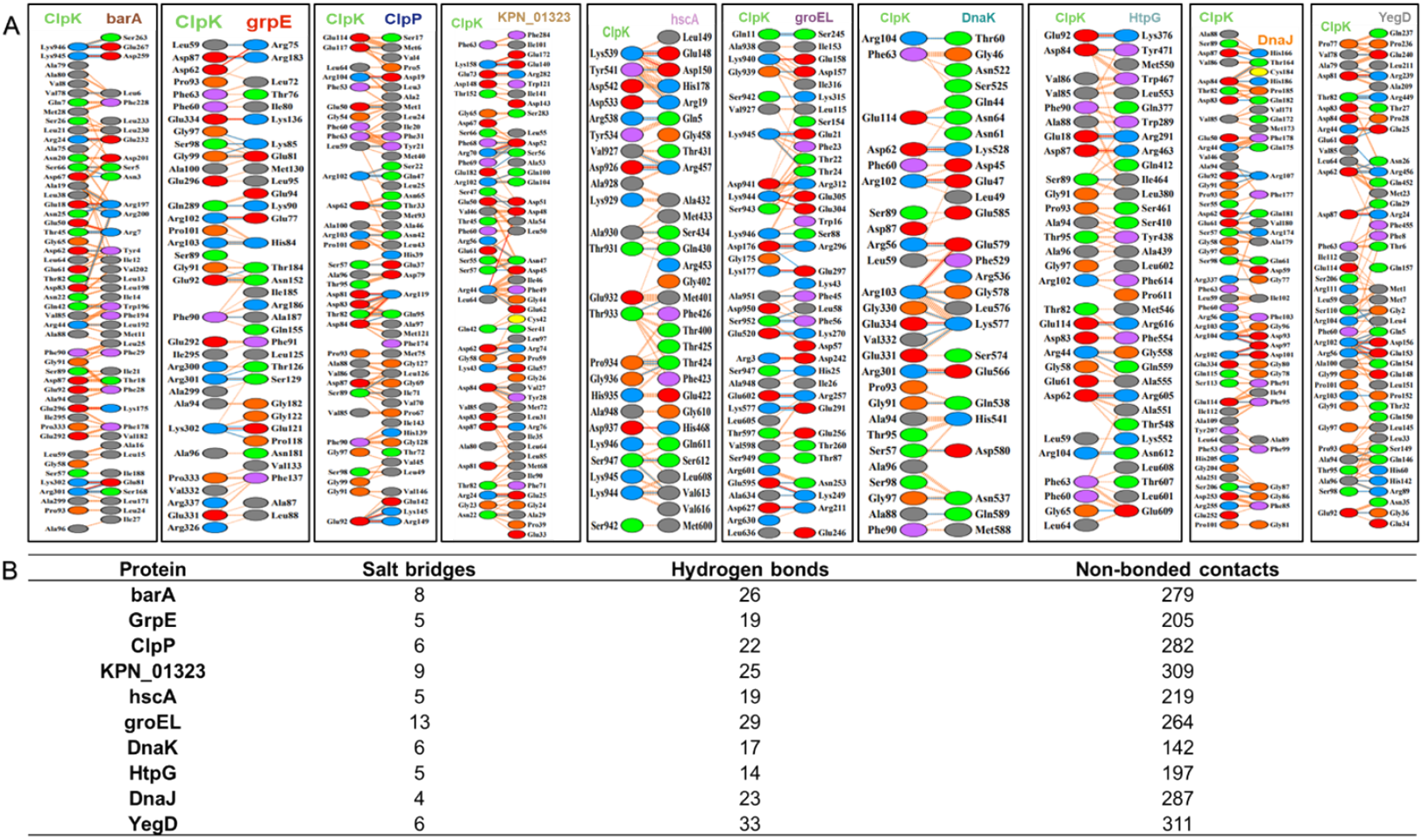
Residue interactions at the interface of ClpK and its putative binding partners. **A)** The interaction diagram is coloured by residue type where red lines represent salt bridges; blue lines represent hydrogen bonds and orange dashes represent non-bonded contacts. **B)** Number of salt bridges, hydrogen bonds and non-bonded contacts for each complex.

The binding affinities and interaction energies between ClpK and its putative protein partners were evaluated based on Gibbs free energy (ΔG) values, dissociation constants (K_d_), and binding free energies. KPN_01323 had the strongest predicted interaction with ClpK as shown by the low ΔG values (-18.6 kcal/mol), and high K_d_ (7.5e-14 M). This correlates with the binding free energy scores observed in Table 1 and the interactions observed in Figure 4. In contrast, DnaK had the lowest ΔG values (12.4 kcal/mol) and K_d_ (1.8e-9 M), indicating relatively lower binding stability. These findings suggest that ClpK may interact with specific protein partners, possibly reflecting functional relevance in protein quality control or stress response pathways.

### 1.2.4. MD simulation analysis of ClpK complexes

RMSD analyses over the 200 ns MD simulation was used to evaluate the conformational stability of both the unbound (*apo*) proteins and their ClpK-bound complexes. Overall, the *apo* forms displayed greater structural fluctuations compared to the bound states (Figure 5). In particular, greater fluctuations in the *apo* state were observed for barA (32.16 ± 12.47 Å) and grpE (24.37 ± 12.47 Å) indicative of significant structural flexibility in their unbound conformations (Figure 5A). In contrast, complex formation with ClpK resulted in reduced RMSD fluctuations (Figure 5B). This suggests that complex formation with ClpK may contribute to enhanced structural stability, except for DnaK where the fluctuations are indicative of increased structural flexibility.

**Figure 5:**
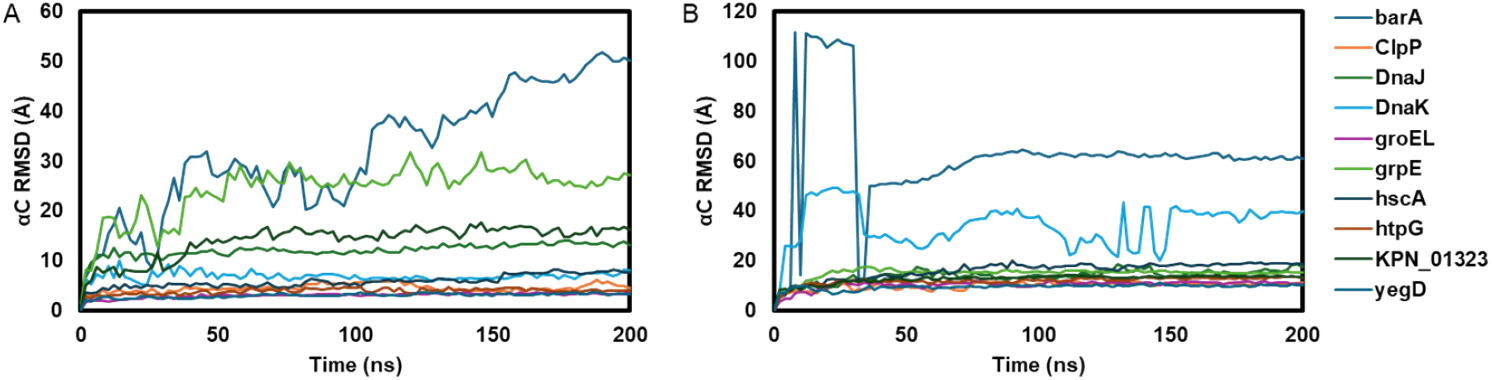
RMSD analysis profiles of alpha-carbon (^α^C) atoms of the putative partners. **A)** RMSD trajectories of *apo* forms. **B)** RMSD trajectories of the ClpK-partner complexes over 200 ns.

To evaluate the overall compactness and structural integrity of the putative ClpKinteracting partners, the radius of gyration (Rg) was calculated over the 200 ns MD simulation (Figure 6). In the *apo* state, the Rg trajectories remained relatively stable with only minor fluctuations, suggesting that the proteins maintained consistent folding and compactness throughout the simulation (Figure 6A). Similarly, the ClpK-bound complexes exhibited comparable Rg stability (Figure 6B), indicating that ClpK binding did not induce major conformational expansion or unfolding (Motiwala *et al*., 2025). Interestingly, a slight increase in Rg was observed for all the complexes except barA, in comparison to their *apo* counterparts. For barA (95.52 ± 0.20 Å), a slight decrease was observed in comparison to the apo form (105.84 ± 0.31 Å). This indicates subtle conformational adjustments to accommodate binding. None of the trajectories exhibited drastic deviations or signs of structural destabilization. Overall, this analysis indicates that proteins retain their compact structure upon ClpK interaction, supporting the stability and integrity of the predicted complexes.

**Figure 6:**
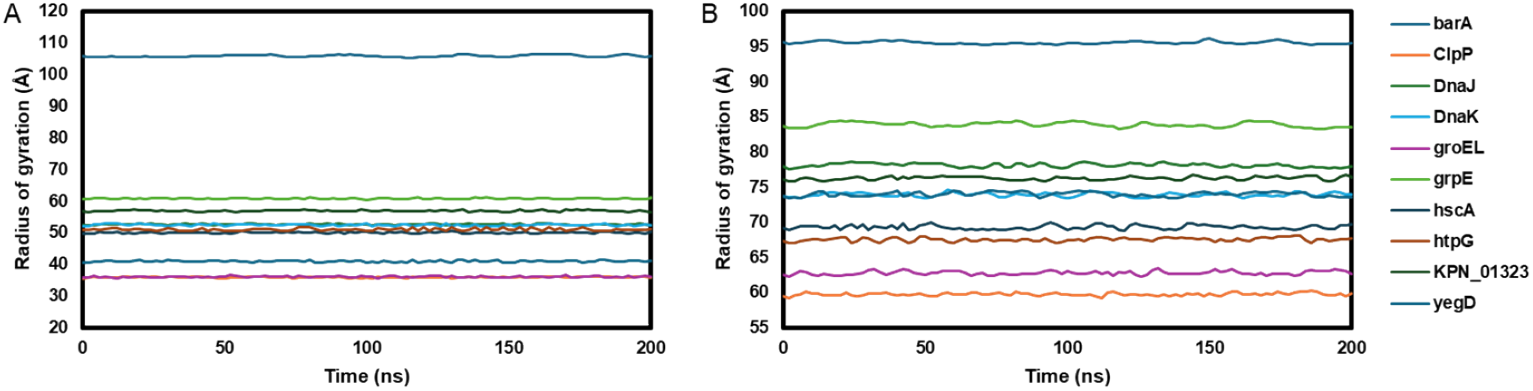
Radius of gyration (Rg) analysis of the putative partners. **A)** Rg trajectories of the *apo* proteins over 200 ns. **B)** Rg trajectories of ClpK-partner complexes.

## 1.3. Discussion

ClpK is a thermotolerant ATPase found across various *Klebsiella* species and has been associated with enhanced heat resistance and improved survival under stressful conditions (Bojer *et al*., 2013, Motiwala *et al*., 2021). Despite its role in thermotolerance and protein quality control, ClpK remains poorly characterized, and therefore there is limited evidence pertaining to its protein-protein interactions (PPIs). It is important to understand the significance of biological and molecular processes of PPIs within cells for identifying novel antimicrobial targets (Shahid *et al*., 2021, Murakami *et al*., 2017). This study represents a computational effort to predict and characterise potential ClpK interaction partners, combining network-based inference, molecular docking and molecular dynamics simulations to investigate the nature and stability of these associations.

The STRING database was used to derive interaction networks using ClpB as a proxy for ClpK (Figure 1), owing to their structural and functional similarity, an approach supported by previous structural and functional comparisons (Motiwala *et al*., 2021). The resulting high-confidence network included ClpP, BarA, KPN_01323, GroEL, GrpE, DnaK, HscA, DnaJ, HtpG and YegD as potential ClpK interaction partners. These proteins are known to participate in proteostasis, stress adaptation, and signal regulation pathways (Straus *et al*., 1990, Susin *et al*., 2006, Lupoli *et al*., 2018, Mangla *et al*., 2023). Docking analyses revealed that all ClpK-partner complexes formed energetically favourable and stable interactions mediated by hydrogen bonds, salt bridges and non-bonded contacts (Figure 4), indicating the formation of stable proteinprotein interactions (Donald *et al*., 2011).

Among the predicted interacting partners for ClpK, ClpP is a caseinolytic protease that functions in conjunction with Clp ATPases through IG(F/L), a conserved interaction motif, to facilitate the degradation of damaged, misfolded, or regulatory proteins (Aljghami *et al*., 2022, Maurizi and Xia, 2004, Queraltó *et al*., 2023). The inclusion of ClpP within the predicted network supports earlier evidence that ClpK contributes to thermotolerance via proteolytic regulation (Bojer *et al*., 2013). Structural modelling revealed that ClpK did not interact with this motif directly, suggesting a non-canonical interaction mechanism distinct from other Clp ATPases. This divergence could reflect evolutionary adaptation, allowing ClpK to interact flexibly with ClpP or alternative partners under stress. Experimental validation of this non-canonical mode of interaction through site-directed mutagenesis or co-immunoprecipitation would be valuable to confirm this hypothesis.

Several other predicted partners, namely, GroEL, GrpE, DnaK, DnaJ, HtpG and YegD are molecular chaperones essential for protein folding, disaggregation, and stress response (Thirumalai *et al*., 2020, Harrison, 2003, Calloni *et al*., 2012, Silberg *et al*., 2004, Kolbe Musskopf *et al*., 2018, Garcie *et al*., 2016, Itoh *et al*., 1999). Their association with ClpK reinforces the view that this ATPase may act as a chaperone hub, coordinating multiple arms of the bacterial heat-shock response. The presence of BarA, a highly conserved histidine kinase which regulates the Csr system, suggests a broader regulatory dimension in ClpK function, possibly linking stress sensing to proteolytic activity (Alvarez *et al*., 2021). Interestingly, the inclusion of KPN_01323, a component of the Type VI secretion system contractile sheath (T6SS), was unexpected and points toward a potential role of ClpK in bacterial competition and pathogenesis (Cherrak *et al*., 2019). Interestingly, ClpK exhibited the strongest and most stable predicted interaction with KPN_01323. Binding energy and interaction energy analyses further supported the strength of ClpK-KPN_01323 interaction, with this complex displaying the most favourable energetics among all tested partners (Table 2). Given the novelty and predicted stability of this interaction, further investigation *in vitro* experimental validation to explore the functional relevance of this interaction in *K. pneumoniae* is warranted.

**Table 2:** Summary of ClpK and putative protein partners interaction energetics.

| Protein | $\Delta G$ (kcal/mol)<sup>1</sup> | $K_d$ (M)<sup>1</sup> | Binding free energy (kcal/mol)<sup>2</sup> |
| --- | --- | --- | --- |
| barA | -17.3 | $6.8e-13$ | -170.54 |
| ClpP | -15.8 | $7e-12$ | -174.96 |
| DnaJ | -16.4 | 2.5e-12 | -186.09 |
| DnaK | -12.4 | 1.8e-9 | -39.43 |
| groEL | -12.5 | 1.6e-9 | -84.37 |
| grpE | -13.5 | 2.9e-10 | -96.91 |
| hscA | -12.4 | 1.9e-9 | -62.69 |
| htpG | -14.0 | 1.3e-10 | -72.86 |
| KPN_01323 | -18.6 | 7.5e-14 | -134.25 |
| yegD | -17.7 | 3.4e-13 | -174.21 |

<sup>1</sup>PRODIGY server <sup>2</sup>HawkDock server – MM-GBSA

MD simulations over 200 ns confirmed the stability of the ClpK-partner complexes. RMSD profiles showed that all complexes equilibrated rapidly and maintained structural integrity throughout the simulation period (Figure 5), while Rg analysis demonstrated consistent compactness across both *apo* and bound forms (Figure 6). These results suggest that ClpK–partner binding does not induce large-scale conformational rearrangements or destabilization, which aligns with ClpK’s proposed role as a thermotolerant ATPase capable of maintaining structure under fluctuating conditions (Motiwala *et al*., 2025). The observed stabilization of key complexes, particularly ClpK–KPN_01323 and ClpK–ClpP, supports their potential physiological relevance and provides structural bases for experimental follow-up.

In conclusion, this bioinformatics study proposes the first prediction of PPIs of ClpK with ten putative interacting partners. These findings provide valuable insight into the potential functional network of ClpK, especially its role in stress response and protein homeostasis in *Klebsiella* species, particularly *K. pneumonia*. It is important to further characterise these predicted interactions through experimental validation methods such as enzyme assays, co-immunoprecipitation, or pull-down assays. Overall, this study provides a foundation for further *in vitro* and *in vivo* investigations that will contribute to understanding the biological role of ClpK and its potential as a therapeutic target.

## 1.4. Materials and Methods

### 1.4.1. STRING Interaction Analysis

In the absence of ClpK in the STRING database (Search Tool for the Retrieval of Interacting Genes/Proteins, version 12; https://string-db.org), the closest homolog, ClpB, was used to construct the protein-protein interaction (PPI) network. The interactions were predicted based on combined scores from experimental data, coexpression, neighbourhood, gene fusion, co-occurrence, and text mining. A highconfidence score threshold (>0.7) was applied to filter predicted associations. The resulting network was visualized using STRING’s integrated viewer, and nodes were annotated according to their predicted functional partners.

### 1.4.2. Protein 3D structures

The three-dimensional structure of ClpK was retrieved from the AlphaFold Protein Structure Database, which uses deep learning approaches to generate high-confidence models based on homology and multiple sequence alignments (Jumper *et al*., 2021). The model with the highest predicted Local Distance Difference Test (pLDDT) score was selected for subsequent analysis. The 3D structures of ClpK-interacting proteins were constructed using the SWISS MODEL server. The amino acid sequences of the interacting proteins were retrieved from the Uniprot database and submitted to the SWISS-MODEL workspace for comparative modelling. The most suitable templates were selected based on sequence identity, GMQE (Global Model Quality Estimation), and Qualitative Model Energy ANalysis (QMEAN) scoring functions. Only models with high structural confidence and favourable stereochemical properties were used for molecular docking and interaction analyses.

### 1.4.3. Protein-protein docking and analysis

Molecular docking of ClpK and its putative interacting partners was performed using the ClusPro 2.0 server. Blind docking was performed without prior specification of the binding interface. The cluster with the highest number of members was selected for further analysis as the most probable binding conformation. The PDBSum server was used to analyse protein-protein interactions (Laskowski *et al*., 2018). The HawkDock server was used to perform MM/GBSA free energy calculations to providing an estimate of the binding stability of the docked protein–protein complex (Zhang *et al*., 2025). These analyses were further complemented using the PRODIGY server to predict the binding affinity (ΔG) of the protein-protein interactions and help validate biological interaction plausibility (Xue *et al*., 2016).

### 1.4.4. Molecular dynamics simulation and post dynamics analysis

Molecular dynamics (MD) simulations were performed on a Linux (Ubuntu) workstation using the GPU-enabled Desmond simulation engine integrated within Maestro v12.2. Prior to simulation, each complex was prepared using the System Builder module in Desmond. This module solvated the systems using the TIP3P explicit solvent model and the OPLS_2005 force field. The complexes were embedded in an orthorhombic box, with a 10 Å buffer between the outermost atom of the solute and box edges, and box angles set at 90^0^. Counter ions were added to neutralize the system, and 0.15 M NaCl was included to mimic physiological strength. The solvated system was then energy minimized before proceeding to the simulation phase. The MD protocol consists of eight stages, with stages 1–7 serving as equilibration and stage 8 as the production run. In stage 1, the system topology and parameters were initialized. Stages 2 and 3 involved Brownian dynamic simulations under NVT conditions at 10 K, lasting 100 ps and 12 ps, respectively, with restraints applied to the solute’s heavy atoms. Pocket solvation (stage 5) was skipped. Stages 4, 6 and 7 were conducted under NPT conditions at 10 K, lasting 12, 12 and 24 ps, respectively. Restraints on heavy atoms were maintained in stages 4 and 6 but removed in stage 7. Finally, stage 8 comprised the production phase, during which as unrestrained 200 ns MD simulation was performed at a constant temperature. Trajectories derived from the MD simulation studies were used for post dynamic analysis using Schrodinger Maestro v12.2. The simulation interaction diagram algorithm was used to analyze the root mean square deviation (RMSD) of the alpha carbon atoms (^α^C) of the complexed and *apo* proteins. The simulation events analysis algorithm was used to analyze the radius of gyration (Rg) of the complex and *apo* proteins.

## 1.5. Author Contributions

Conceptualization, T.K., and I.A.; methodology, T.B., and T.K.; data curation T.B.; modelling, T.B., and T.K.; validation, T.B., and T.K.; molecular dynamic simulations, T.B.; formal analysis, T.B. and T.K.; investigation, T.B. and T.K.; protein expression and optimization, T.B.; biophysical characterization, T.B., and T.K. writing—original draft preparation, T.B.; writing—review and editing, T.B., I.A., and T.K.; visualization, T.B., and T.K.; supervision, T.K., and I.A.; project administration, T.K., and I.A.; All authors have read and agreed to the published version of the manuscript.

## 1.6. Funding

Tehrim Ballim would like to thank Department of Science and Technology-National Research Foundation (DST-NRF), South Africa for the Doctoral Scholarship with Grant Number MND210602605517. Thandeka Khoza also thanks the National Research Foundation Grant number 121275, South African Medical Research Council-SIR grant and University of KwaZulu-Natal for research grants.

## 1.7. Conflicts of Interest

The authors declare no conflict of interest. The funders had no role in the design of the study; in the collection, analyses, or interpretation of data; in the writing of the manuscript, or in the decision to publish the result.

## Notes

### Competing Interest Statement

The authors have declared no competing interest.

